# Arbuscular mycorrhizal fungal composition across US citrus orchards, management strategies, and disease severity spectrum

**DOI:** 10.1101/2022.03.01.482593

**Authors:** MengYuan Xi, Elizabeth Deyett, Nichole Ginnan, Vanessa E.T.M. Ashworth, Tyler Dang, Sohrab Bodaghi, Georgios Vidalakis, M. Caroline Roper, Sydney I. Glassman, Philippe E. Rolshausen

## Abstract

Arbuscular mycorrhizal fungi (AMF) remain understudied in perennial cropping systems. Citrus is a globally grown fruit tree and under threat by the pandemic Huanglongbing (HLB) disease. Here, we assessed in what capacity geographical location, management strategies and disease affect AMF citrus root communities. Root samples were collected from 88 trees in ten orchards located in the two major citrus producing states in the US. Orchards were selected based on conventional or organic practices in California and based on HLB symptom severity in Florida. We used AMF-specific amplicon sequencing primers to capture community composition and diversity. Taxa names were assigned based on a phylogenetic analysis that comprised a backbone of AMF references sequences from Mycobank and virtual taxa from the MaarjAM database. AMF were detected in 78% of citrus root samples with taxa belonging to six known (*Dominikia, Funneliformis, Glomus, Rhizophagus, Sclerocystis, Septoglomus*) and unknown Glomeraceae genera. Geographical location affected AMF community composition but not richness, whereas management practice and disease influenced both richness and composition. Our approach indicated that perennial agroecosystems share a set of AMF generalist and specialist taxa. Some taxa could improve environmental fitness and be exploited for agricultural purposes.

## Introduction

Arbuscular mycorrhizal fungi (AMF) are biotrophic organisms forming symbiotic associations with more than 70% of the land plants across a broad range of terrestrial ecosystems (1, 2). The nature of the mutualistic symbiosis resides in a trade-off between a more efficient root acquisition of water and nutrients (especially phosphorus) via the mycorrhizal hyphal network, in exchange for photo-assimilated carbon. The outcome of this interaction often results in improved plant environmental fitness with increased tolerance to biotic and abiotic stresses (3, 4). In addition, AMF improve soil structure by forming stable soil aggregates thereby limiting erosion and leaching of nutrients (4, 5).

Plant parts above and below ground are intricately connected and the health status of the root system often determines plant growth and productivity. The rhizosphere microbial diversity is a biomarker of soil fertility and plays a central role in sustainable agricultural systems (6, 7). Low input agriculture systems (organic and biodynamic farming) have adopted cultural practices to rely on soil biological metabolism and function to support soil fertility and plant root health. In contrast, intensive farming practices characterized by monoculture, input of synthetic agrochemicals, and/or soil disturbance caused by mechanical stress generally leads to degradation of soil ecosystem biodiversity. Mycorrhizal fungi are major participants of the rhizosphere microbiome and are under the constraints of farming practices, with research indicating that mycorrhizal fungi are more diverse under organic agroecosystems (8, 9). Understanding the factors that shape AMF communities could improve management tools and recommendations for implementing more sustainable agricultural practices.

Citrus is a high-value crop and one of the most popular and widely grown fruit trees globally. It is praised for its nutritional values and benefits to human health as a source of vitamins, fibers, and minerals. Citrus accounts for 16% of the total value of U.S. fruit production (10) with California and Florida leading the nation’s fresh fruit and juice markets, respectively. Symbiotic associations with AMF have been reported in all citriculture production areas and AMF communities are shaped by edaphic characteristics, orchard management practices, and host variety and age (11–14). Adopting low input farming practices for citrus at a large geographic scale has been challenging because of Huanglongbing (HLB), a disease associated with an invasive phloem-limited bacteria in the *Candidatus* Liberibacter genus (i.e., *C*. L. *asiaticus, americanus*, and *africanus*) (15). In the U.S., *C*. L. *asiaticus* is the primary threat to citrus. The Florida citrus industry suffered a 74% decline in production with losses amounting to over $1 billion annually because of HLB (10, 16). The pathogen is vectored by an invasive insect (*Diaphorina citri*; the Asian citrus psyllid) and disease management has been mostly achieved by intensive regimens of synthetic insecticide applications to control insect populations (17). In heavily affected orchards, trees are also treated with antibiotics (oxytetracycline and streptomycin) to reduce levels of pathogen inoculum reservoirs (18). Those practices have raised environmental concerns due to the risk of unintended consequences for biodiversity and selection for resistance in bacterial and insect populations (19, 20).

In HLB-impacted orchards, the tree rhizosphere suffers from microbial dysbiosis and root collapse, thereby weakening the host and its defense response against attack from other pathogens (21, 22). Specifically, a study from Florida found that high relative abundance of Glomeromycota correlated with healthier trees (22), although the ITS2 amplicon used in the study limited the resolution of AMF taxonomy and may have omitted key taxa (23). AMF have been established to provide protection against citrus root diseases (24, 25), and developing practices that support their biodiversity can offer new ground for management strategies. Previous studies have found at least seven genera of AMF associated with citrus roots (*Acaulospora, Entrophospora, Gigaspora, Glomus, Pacispora, Sclerocystis*, and *Scutellospora*), yet little is known about how these communities vary across geographies or management practices. Moreover, many of these studies are morphological-based and may have underrepresented AMF diversity that can be captured with sequencing approaches (26).

Citrus is an emblematic specialty crop to US agriculture. In the wake of the economic and environmental challenges posed by HLB, alternative strategies to farming citrus should be considered. This study aimed to profile AMF community composition and structure in citrus agroecosystems from two distinct climatic zones within the continental US. We captured citrus root-associated AMF diversity by employing AMF-specific amplicon sequencing primers (27, 28) to better identify taxa and capture the biodiversity associated with citrus roots. Orchards in California were selected according to the farming practices (organic vs. conventional) and in Florida according to severity of HLB disease symptoms expression (mild, moderate, severe) because HLB positive trees have not been detected in commercial groves in California. We hypothesize that AMF community composition will overall differ between California and Florida because temperature and pH are known to be major drivers of AMF composition globally and these regions have inherently different climatic conditions and edaphic factors (29). We also anticipate some level of overlap in AMF taxonomic distribution between these two citriculture areas because of the similarity of the host. In addition, within each citrus area, we speculate that organic farming will have higher AMF diversity than conventional farming and that HLB affected trees will display fewer AMF symbionts because of the lower photoassimilated carbon transfer to the host root system. This study provides insightful information about AMF dynamics within a major perennial agroecosystem and identifies putative generalist and specialist taxa that could be exploited for agricultural purposes.

## Materials and Methods

### Root sample collection

Root samples were collected from 88 trees in ten citrus orchards located in Florida (ten trees per orchard) and California (eight trees per orchard) following published protocols (22). In Florida, 40 samples were collected in March 2017 from four conventional orchards (Table 1). All trees were rated for HLB symptoms using a disease rating scale ranging from mildly, to moderately and severely symptomatic. In California, 48 samples were collected in October 2017 from four conventional and two organic orchards (Table 1). For the California sampling, feeder roots were sampled from two sides of the tree approximately 0.5 m away from the base of the trunk and sealed in a plastic bag. Gloves were changed and clippers and shovels were sterilized with 30% household bleach between each sampled tree. All samples were immediately placed on ice in a cooler for transit to the laboratory and were frozen for shipment to the University of California, Riverside (UCR). Root samples were rinsed with autoclaved purified water (Barnstead Mega-Pure System MP-6a, Thermo Fisher Scientific, Waltham, MA, USA) and approximately 5 g of rinsed root tissue was placed into 50 ml conical tubes, stored at −80°C, and then lyophilized (Labconco FreeZone 4.5L, Kansas City, MO) for 16 to 20 hours. Samples were collected similarly in Florida and lyophilized prior to shipment to UCR under USDA permit #P526P-16-00352 on dry ice.

**Table 1:** Location and farming practices for the citrus orchards sampled in this study.

| Orchard # | US State | Number of Root Samples Collected | Orchard Type | Scion Variety | Rootstock | Number of Root samples with AMF |
| --- | --- | --- | --- | --- | --- | --- |
| 1 | Florida | 10 | Conventional | Valencia | Swingle | 9 |
| 2 | Florida | 10 | Conventional | Ruby Red | US897 | 9 |
| 3 | Florida | 10 | Conventional | Parson Brown | Swingle | 9 |
| 4 | Florida | 10 | Conventional | Hamlin | Sour Orange | 6 |
| 5 | California | 8 | Conventional | Newhall x Satsuma | Carrizo | 7 |
| 6 | California | 8 | Conventional | Late Navel Powell | Carrizo | 8 |
| 7 | California | 8 | Conventional | Late Navel Powell | Carrizo | 2 |
| 8 | California | 8 | Conventional | Late Navel Powell | Carrizo | 4 |
| 9 | California | 8 | Organic | Newhall x Satsuma | Carrizo | 7 |
| 10 | California | 8 | Organic | Late Navel Powell | Carrizo | 8 |

### DNA extraction, library construction and sequencing

DNA was extracted from roots according to published protocols (22). Frozen and freeze-dried roots were crushed into small pieces (<0.5 cm) with sterile stainless-steel spatulas on dry-ice, and 100 mg of freeze-dried tissue was transferred to 2-ml microcentrifuge tubes (Eppendorf Safe-Lock tubes; Eppendorf, Hamburg, Germany) containing a single 4-mm stainless-steel grinding ball (SPEX SamplePrep, Metuchen, NJ, U.S.A.). Samples were chilled at −80°C for 15 min, then pulverized to a powder using a 2010 Geno/Grinder (SPEX SamplePrep) at 1,680 rpm for 20 to 30 seconds, twice. Then, 1 ml of 4 M guanidine thiocyanate buffer was added to the pulverized root samples. Samples were incubated at 4°C for 15 min and subsequently centrifuged for 1 h at 17,500 × g. DNA was isolated using the MagMAX-96 DNA Multi-Sample Kit (Thermo Fisher Scientific) with the protocol “4413021ForPlants” on a MagMAX Express-96 Deep Well Magnetic Particle Processor. The final DNA was eluted in 100 ml of DNA elution buffer and stored at −20°C prior to Illumina library construction.

DNA was PCR-amplified as described in Phillips et al. (30) targeting the 18S region using the Glomeromycotina-specific AML2 and the universal eukaryote WANDA primer sets (27, 28). Library construction was conducted in a two-step procedure (31) in which first-round amplifications were carried out with primers possessing universal tails synthesized 5′ to the locus-specific sequences (32) and second round-amplifications ligated Illumina MiSeq flowcell adapters and barcodes (30). Recipe for PCR1 included 1 μl of template DNA, 12.5 μl AccuStart II PCR ToughMix (2X) (Quantabio, Beverly, MA), 0.5 μl of each primer (10 μM), and 10.5 μl nuclease-free water, resulting in a 25 μl reaction. Thermocycler conditions for PCR1 were as follows: 2 min at 94°C; 29 cycles of 30 s at 94°C, 30 s at 60°C, 45 s at 68°C. Reaction products were verified on a 1% agarose gel and purified using the AMPure XP magnetic Bead protocol (Beckman Coulter Inc., Brea, CA, USA). PCR2 was performed in a 25 μl reaction, with 1 μl of the undiluted purified PCR1 product, 6.5 μl AccuStart II PCR ToughMix (2X) (Quantabio, Beverly, MA), 2.5 μl of each barcode primer (1 μM), and 12.5 μl nuclease-free water. Thermocycler conditions for PCR 2 were as follows: 2 min at 94°C; 9 cycles of 30 s at 94°C, 30 s at 60°C, 1 min at 72°C. We checked indexed PCR products on an agarose gel and pooled the products by band strength as in Glassman et al. (33) with 1 μl for strong bands, 2 μl for medium bands, and 3 μl for weak bands prior to AMPure bead purification. The purified library was quantified with a Qubit 4 Fluorometer (Thermo Fisher Scientific, Waltham, MA) and quality checked with an Agilent BioAnalyzer 2100 for size and concentration and sequenced with Illumina MiSeq nano run (2 x 250 bp) at the UC Riverside Institute for Integrative Genome Biology. Sequences were submitted to the National Center for Biotechnology Information Sequence Read Archive under accession number SRPXXX.

### Bioinformatics and taxonomy assignment

Initial quality filtering of sequences was done using Trimmomatic (34) truncating reads once the average quality of 5 consecutive base pairs dropped below a quality score of 20. Using DADA2 (v 1.14.1), reads were processed further to remove reads with more than one N calls, ambiguous calls, reads identified as PhiX, reads that were too short, and reads with more than 2 expected errors per DADA2’s algorithm. DADA2 was also used to dereplicate, learn error rates, and create an amplicon single variant (ASV) sequence table (35). Samples with less than 1,000 reads were removed, as were taxa not identified as fungi via the NCBI database. Rare taxa, defined as taxa which were prevalent in less than 2% of samples were also removed. This additional filtering step resulted in 131 total ASVs and 1,085,960 reads. Taxonomy was assigned using BLASTN and the MaarjAM database (36)(downloaded on April 1, 2020) using a cut-off e-value of 1e-50 and assigning virtual taxa (VT) based on lowest e-value and best-annotated sample. MaarjAM is an AMF curated database that is standardized, and comparable across research projects, and preserved in time (23). MaarjAM clustered the 131 ASVs into 32 unique VT, and with 1 sequence left unidentified by the MaarjAM database using the parameters mentioned above. Genus names were assigned through the curation of a maximum likelihood bootstrap tree using the Krüger et al. sequences (37). The 33 representative sequences were 220 bp in length and were aligned in Muscle v.3.7 (38) using default parameters to the 58 consensus sequences from Krüger et al. (37) and updated by Stefani et al. (26). Twelve sequences from GenBank were added, resulting in a total of 103 sequences and an alignment length of 756 nucleotides, consisting of 508 conserved, 235 variable, 176 parsimony-informative, and 57 singleton sites. A maximum likelihood tree was constructed in RAxML v8.2.12 (39) implemented on the CIPRES Gateway (40) using a GTRGAMMA evolutionary model of nucleotide substitution and with branch support inferred using 1000 bootstraps. The tree was rooted using *Paraglomus occultum* as the outgroup as per Krüger et al. (37). The consensus tree was visualized and annotated in iTOL (Interactive Tree of Life; (41)).

### Statistical analyses and data visualization

The R v4.1.1 (R Core Team, 2021) was used to perform statistical analysis and data visualization with the aid of the phyloseq v1.36.0 (42) and ggplot2 v3.3.5 packages (43). Data was transformed using the variance stabilization method in the DESeq package v1.32.0 (44). Alpha diversity was estimated as the number of observed taxa of each sample. Statistical significance was calculated by a generalized linear model using Poisson regression and statistical significance on pairwise comparison was performed through Tukey’s test using the multcomp packages v1.4.17 (45). Beta-diversity plots were created using the Bray-Curtis dissimilarity matrix and NMDS ordination matrix using the Vegan package v2.5.7 (46). Adonis tests with 999 permutations were run to determine statistical differences between categorical variables. For heatmaps, created using ComplexHeatmap v2.9.4 (47), the normalized data were aggregated to the VT level and dataset split based on category. VT occurring in fewer than 3 samples of the dataset were removed for ease of visualization. This resulted in 27 taxa in the heatmap comparing California and Florida samples, 15 taxa in the heatmap comparing management strategy and 17 taxa in the heatmap comparing disease rating. For the Venn diagram, a 10% prevalence filtering was applied to find unique and shared taxa amongst categories. Prevalence was defined as occurring at least 1 time in 10% of the samples within a category. Finally, DESeq2 (V1.32.0; (44)) using the default Wald test and local fit, was used to identify taxa with differential abundance analysis among variables of interest. For this analysis, taxa were aggregated to the VT level. Virtual taxa that had a p-value < 0.01 are represented in the heat maps with a “*” symbol or by colored block.

## Results

We detected AMF in 69 of the total 88 citrus root samples (78%, Table 1) with a total of 131 ASVs and 33 virtual taxa. The maximum likelihood phylogenetic analysis that included several taxa from Krüger et al. (37) and the same outgroup (*Paraglomus occultum*), yielded a similar tree topology with strong bootstrap support at the Family level and indicated that all the VT belong to the Glomeraceae (Fig. 1). Taxonomic identification at the genus level was more challenging for some groups given the short nucleotide sequence length, but monophyletic clades with good bootstrap support were obtained for *Sclerocystis*, *Glomus*, and *Septoglomus*. The genus *Rhizophagus* clustered in a monophyletic clade, but with weak bootstrap support, and only formed a well-supported clade with *Sclerocystis*. Similarly, *Funneliformis*—though itself not a monophyletic group—formed a strongly supported clade with *Septoglomus*. Support for *Dominikia* was low and good bootstrap support was obtained only for a subclade composed of two VTs (VTX00222 and VTX00125) with *Dominikia indica*, and for a subclade composed of *D. iranica* and VTX00155. Based on the tree phylogeny, we assigned twelve VTs (VTX00125, −130, −132, −146, −155, −156, −159, −166, −175, −222, −304 and ‘Unknown’) to *Dominikia*, ten VT (VTX00080, −83, −92, −99, −100, −105, −113, −114, −115, −248) to *Rhizophagus*, four VT (VTX00063, −64, −331, −409) to *Septoglomus*, one VT (VTX00197) to *Glomus*, and one VT (VTX00067) to *Funneliformis*. Placement for five VTs (VTX00075, −214, −301, −323, −384) remained uncertain as they did not cluster in any of those clades and were labeled as Glomeraceae species.

**Figure 1.**
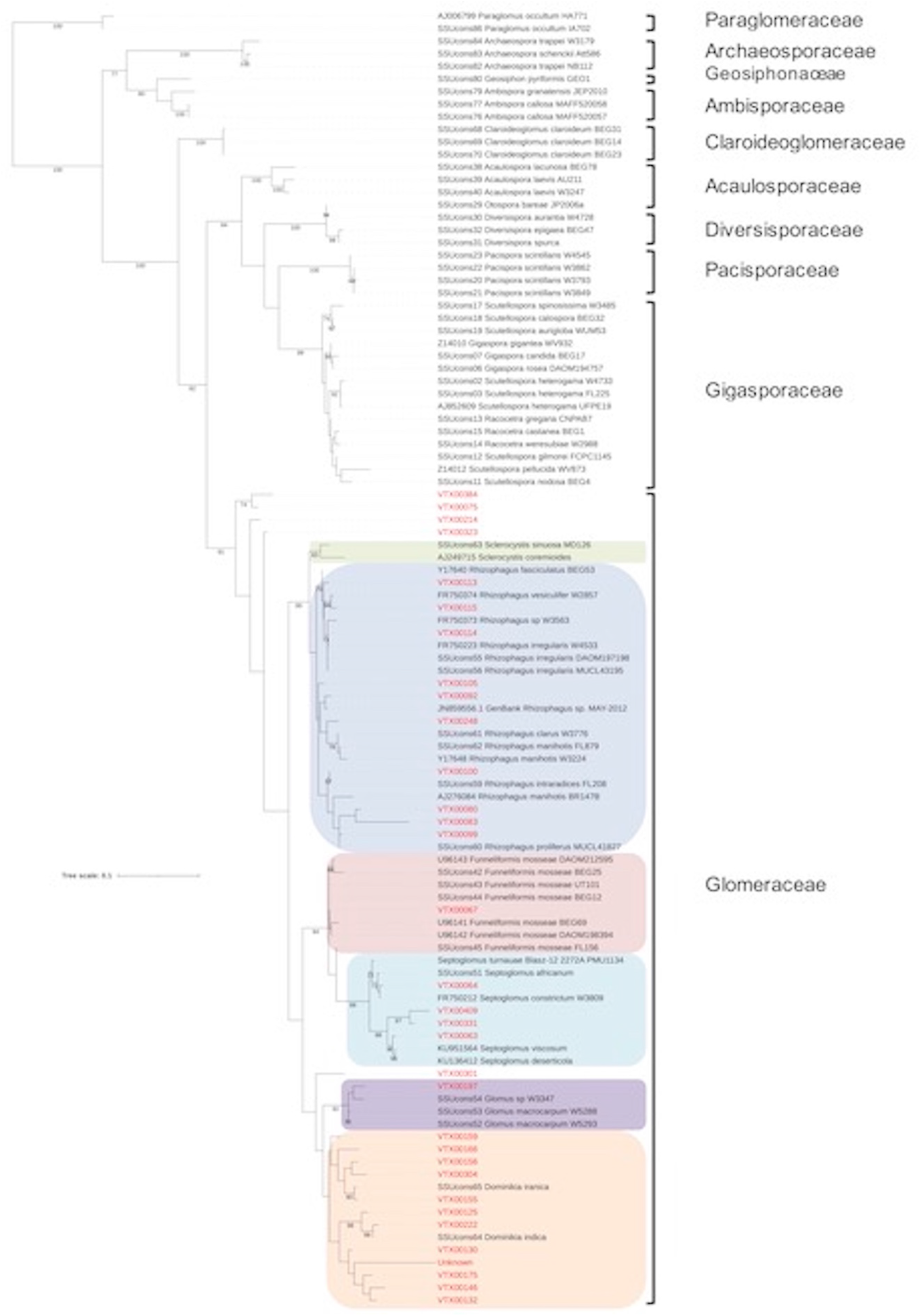
RAxML phylogenetic tree reconstructed by maximum likelihood analysis showing the genus taxonomy assignment of the 33 virtual taxa (in red). The tree represents 18 AMF genera and VT clustered with 5 colored AMF clades. Bootstrap values greater than 70 are displayed on the nodes.

Geographical location did not affect AMF richness in California and Florida citrus orchards as indicated by similar alpha diversity indices (Fig.2A; *P* > 0.05, Poisson generalized linear model with a pairwise Tukey test), although AMF community composition was significantly distinct between the two states (Fig. 3A; Adonis R^2^ = 0.176, *P* < 0.001) with specific taxa and differential abundance among taxa indicated in the heatmap (Fig. 4). Orchard management strategy significantly affected alpha and beta diversity of AMF communities in California, with lower taxa richness in conventional orchards (Fig. 2B; *P* < 0.0001, Poisson generalized linear model with a pairwise Tukey test) and higher compositional variability (i.e., dispersion) among conventional orchard than organically managed orchards (Fig. 3B; Adonis R^2^ = 0.145, *P* < 0.01; betadisper *P* < 0.05). The differential abundance among taxa between organic and conventional farming methods are indicated in the heatmap with compositionally less diverse communities in the conventional orchards (Fig. 5). In Florida, HLB disease status significantly impacted AMF community alpha diversity, with a decline in richness (Fig. 2C; *P* < 0.001, Poisson generalized linear model with a pairwise Tukey test) and shifts in community composition as disease severity increased from mild to severe (Fig. 3C; R^2^ = 0.167, *P* < 0.05; Adonis). The differential abundance among taxa across the disease spectrum are indicated in the heatmap with compositionally less diverse communities in severely HLB-symptomatic trees (Fig. 6)

**Figure 2.**
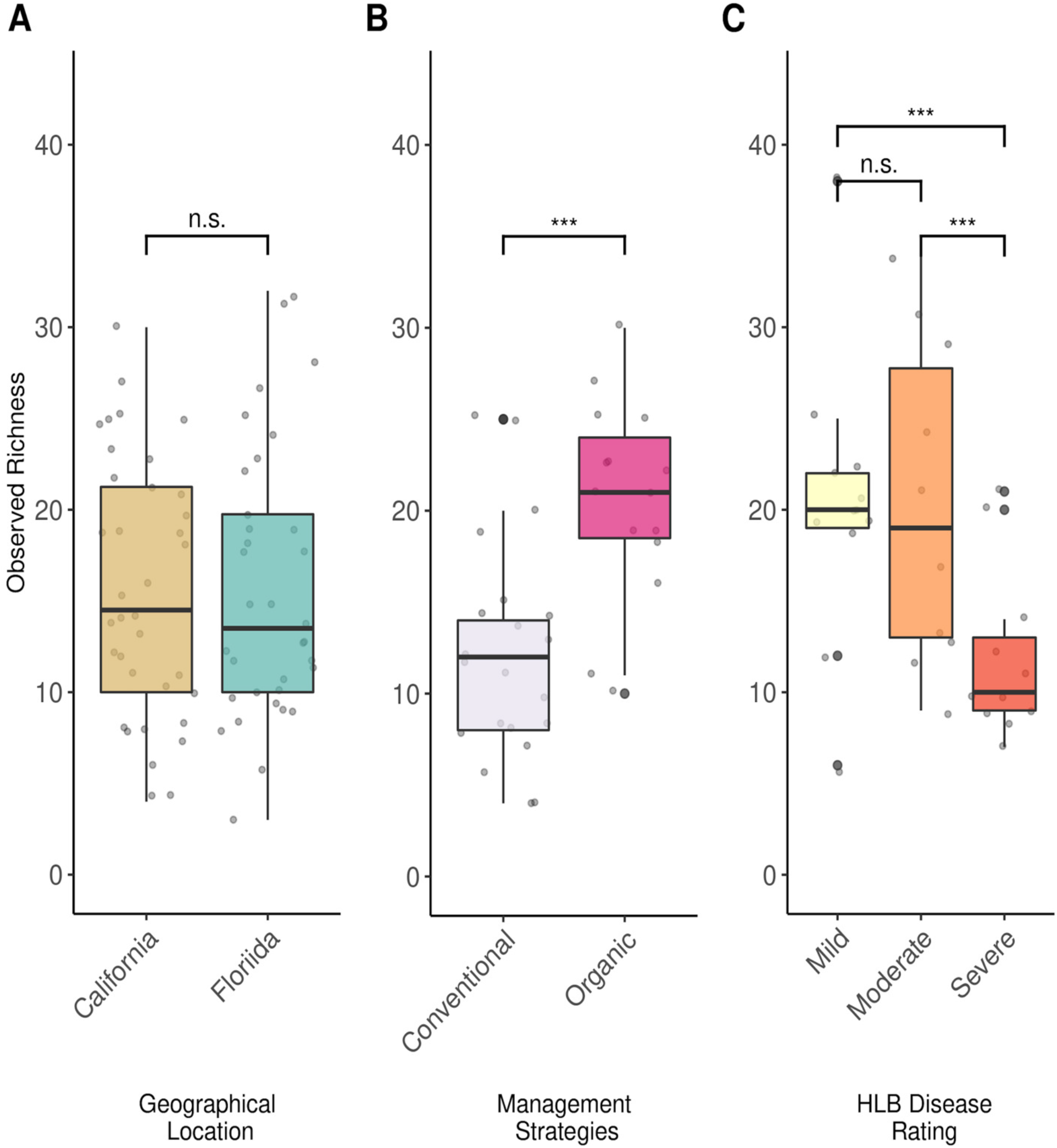
Alpha diversity plots comparing AMF richness across sample types; geographical location (A) shows no effect on richness, unlike management strategies (B) and HLB disease (C). Statistical significance is indicated for *P* < 0.001 (***) based on Poisson generalized linear model with a pairwise Tukey test.

**Figure 3:**
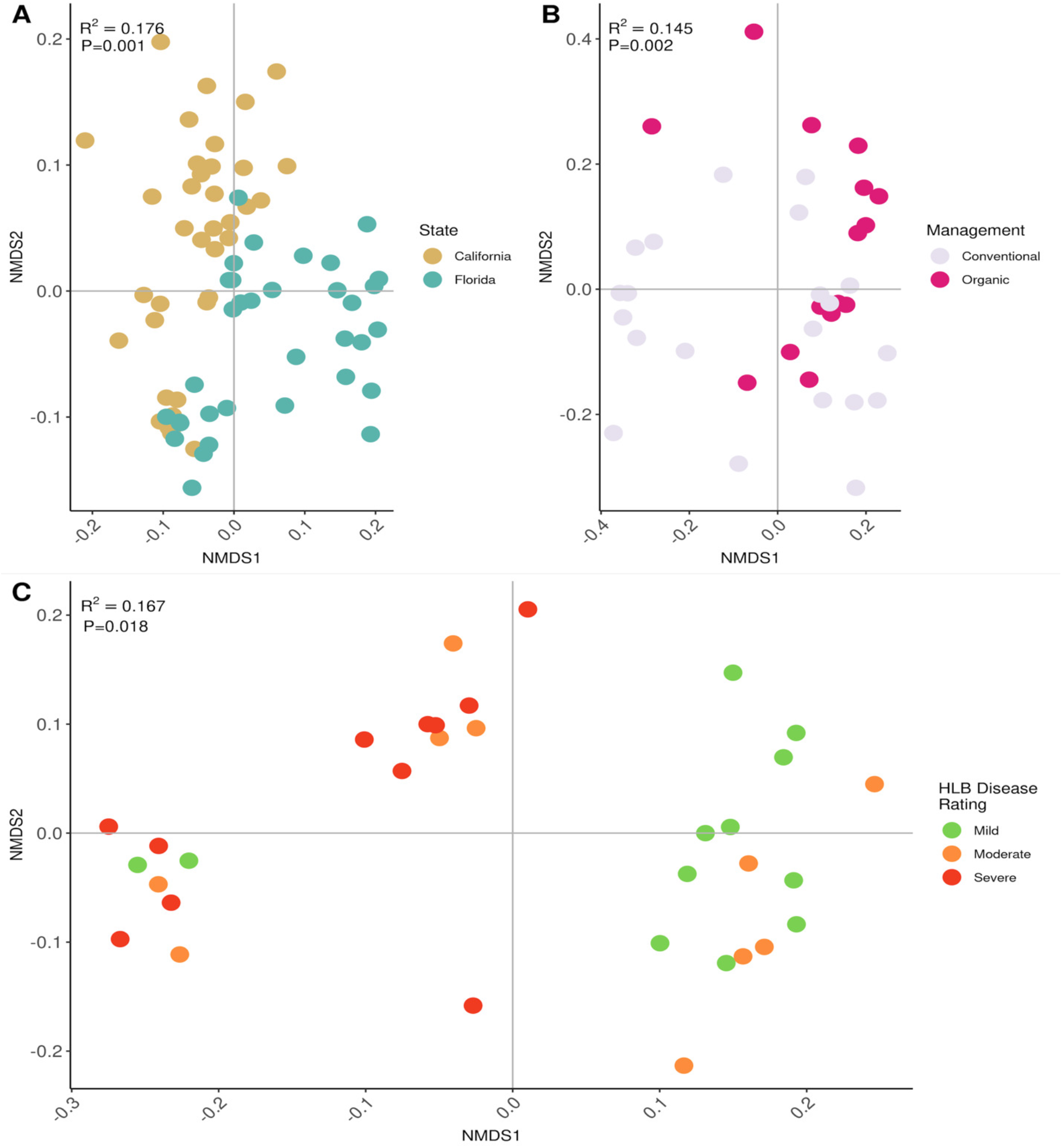
NMDS plots indicating that AMF beta diversity is significantly affected across sample types based on; (A) geographic location (A); management strategies in California (B); and HLB disease in Florida (C). Each dot represents the AMF community composition of a single tree. Points are colored by each group P-values and R^2^ values were measured by permutational multivariate analysis of variance (Adonis) with values shown on the graphs.

**Figure 4:**
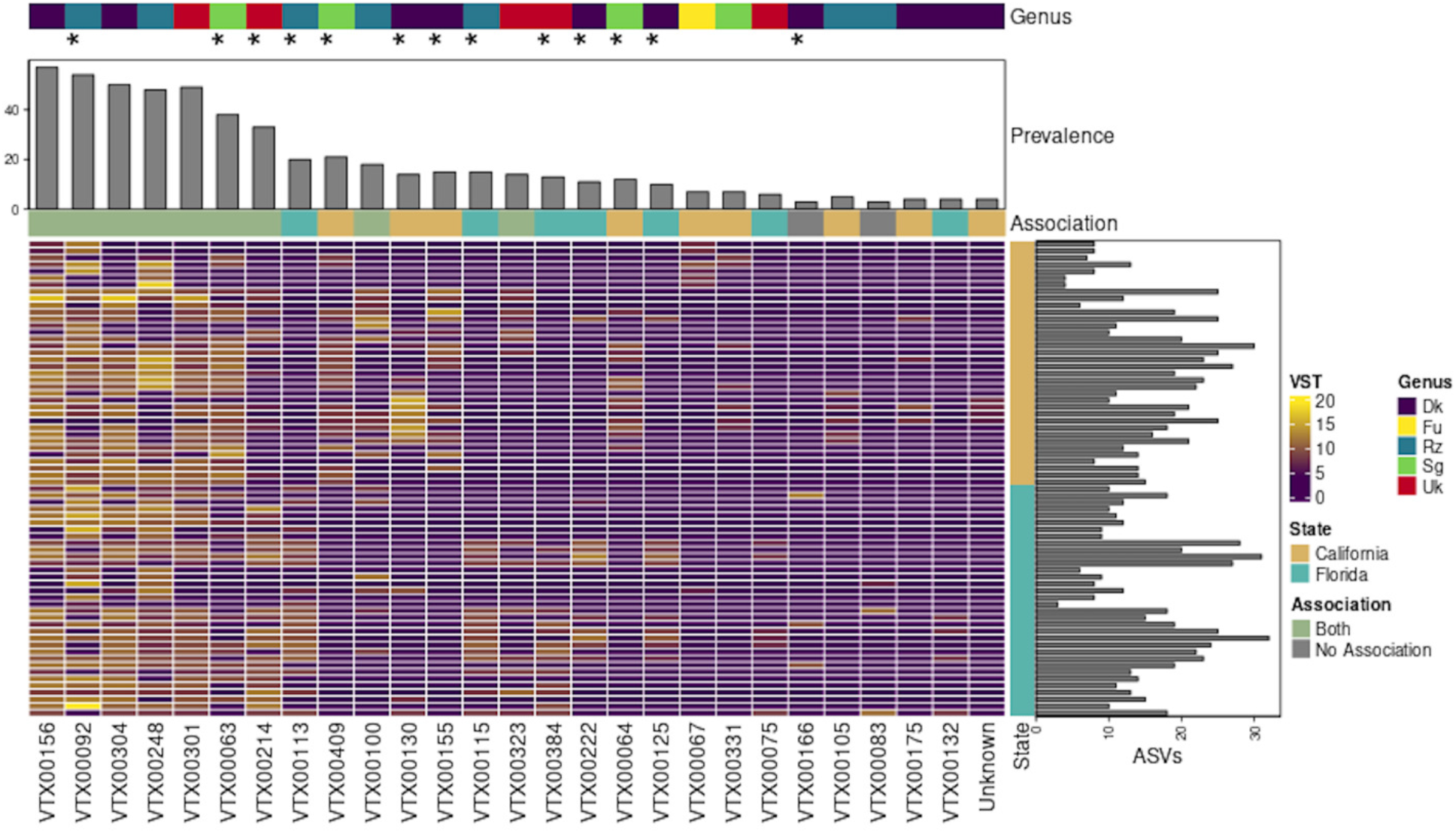
Heatmap of virtual taxa counts between geographical location following Deseq2 variance stabilization transformation. Each row represents a single root sample, and each column represents a unique virtual taxon. For clarity only virtual taxa that occurred in three or more samples are displayed. Top column annotation depicts the genus to which the virtual taxa clustered with based on Maximum likelihood tree in figure 1. Asterisk (*) depict virtual taxa differentially significantly abundant between geographical location per DESeq2 Wald’s test. Column annotation bar graph depicts the prevalence (number of unique samples) the virtual taxa were found in. The “Association” column annotation is a colorimetric Venn diagram with a 10% prevalence cut-off where yellow squares represent taxa associated with California samples, blue squares represent taxa associated with Florida samples and green squares represent taxa which were associated with both Florida and California citrus roots. Gray boxes indicate taxa that did not pass the 10% prevalent cut-off to be associated with either category. Row annotations show which samples belong to each geographical location and bar graph shows the number of unique ASVs associated with each sample. Dk: *Dominikia*; Fu: *Funneliformis*; Rz: *Rhizophagus*; Sg: *Septoglomus*; Uk: Unknown.

**Figure 5.**
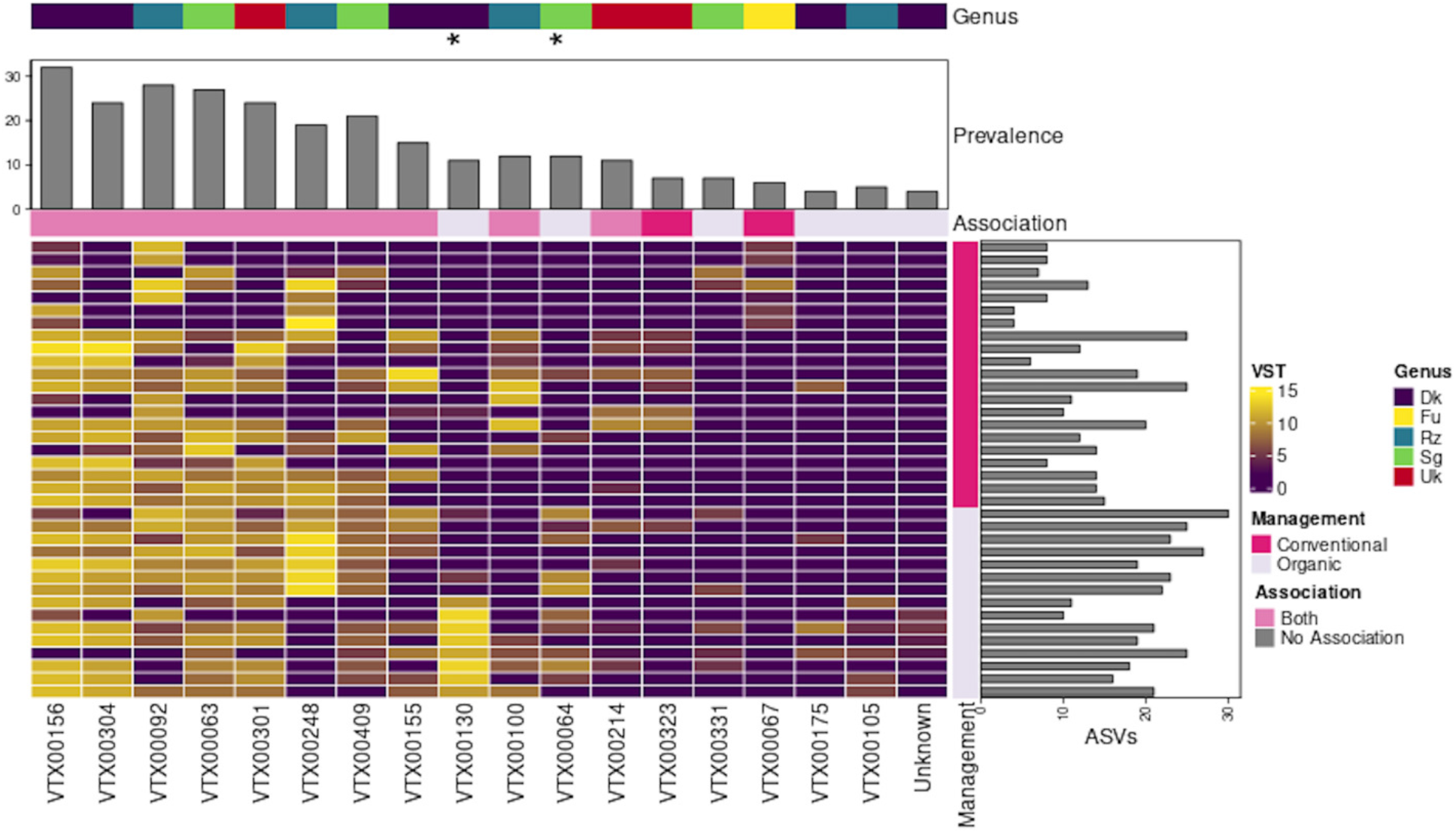
Heatmap of virtual taxa counts between management strategy from California samples following Deseq2 variance stabilization transformation. Each row represents a single root sample, and each column represents a unique virtual taxon. For clarity only virtual taxa that occurred in three or more samples are displayed. Top column annotation depicts the genus to which the virtual taxa clustered with based on Maximum likelihood tree in figure 1. Asterisk (*) depict virtual taxa differentially abundant between geographical location per DESeq2 Wald’s test. Column annotation bar graph depicts the prevalence (number of unique samples) the virtual taxa were found in. The “Association” column annotation is a colorimetric Venn diagram with a 10% prevalence cut-off where dark pink squares represent taxa associated with Conventional samples, lavender squares represent taxa associated with Organic samples and medium pink squares represent taxa which were associated with both Organic and Conventional citrus roots. Row annotations show which samples belong to which management strategy and bar graph shows the number of unique ASVs associated with each sample. Dk: *Dominikia*; Fu: *Funneliformis*; Rz: *Rhizophagus*; Sg: *Septoglomus*; Uk: Unknown.

**Figure 6.**
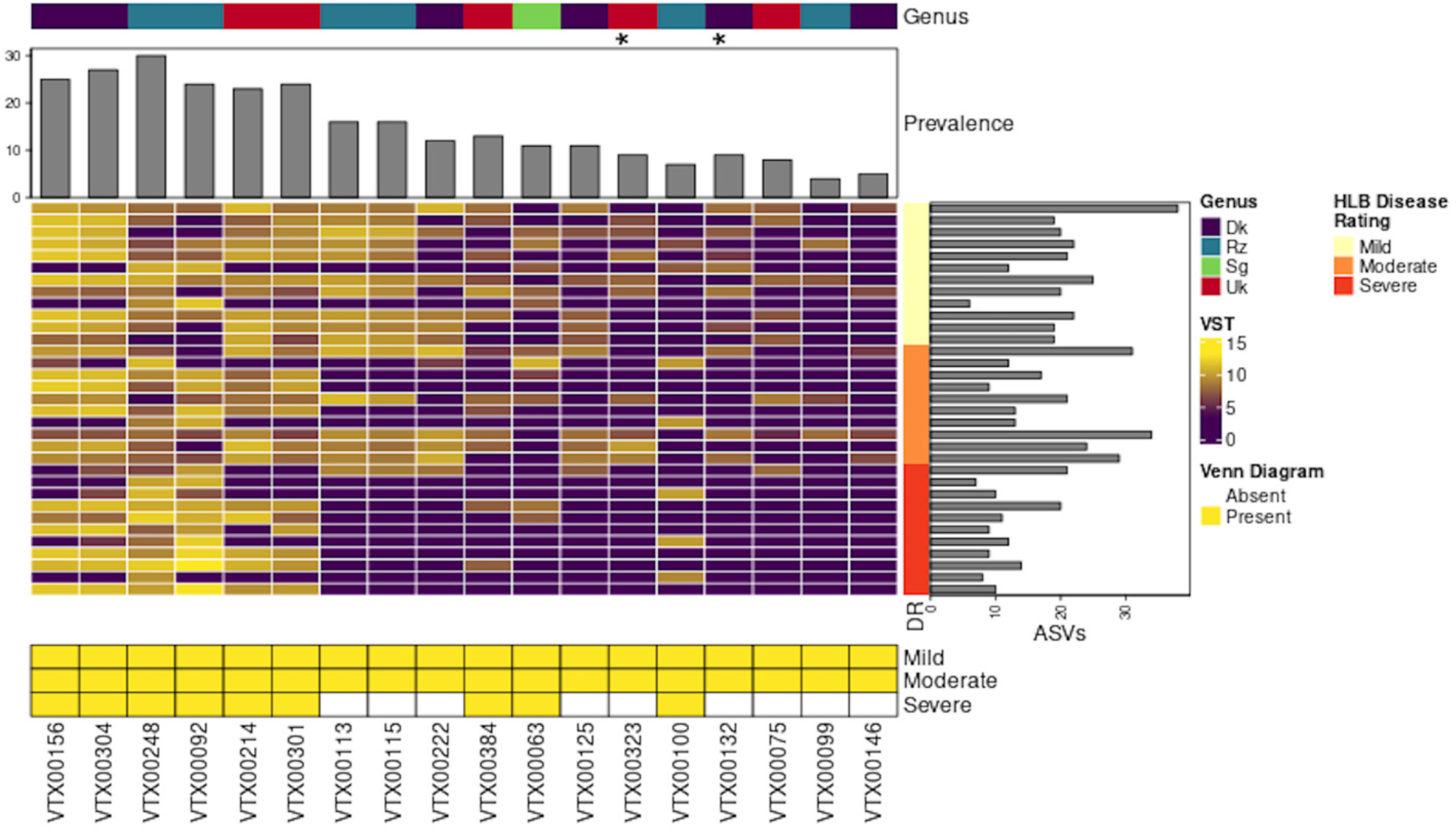
Heatmap of virtual taxa counts of HLB disease severity from Florida samples following Deseq2 variance stabilization transformation. Each row represents a single root sample, and each column represents a unique virtual taxon. For clarity only virtual taxa that occurred in three or more samples are displayed. Top column annotation depicts the genus to which the virtual taxa clustered with based on Maximum likelihood tree in figure 1. Asterisk (*) depict virtual taxa differentially abundant between geographical location per DESeq2 Wald’s test. Column annotation bar graph depicts the prevalence (number of unique samples) the virtual taxa were found in. Row annotations show which samples belong to each disease rating severity and bar graph shows the number of unique ASVs associated with each sample. Venn diagram heatmap with yellow rectangles show which taxa were found in at least 10% of sample from each disease severity. Dk: *Dominikia*; Rz: *Rhizophagus*; Sg: *Septoglomus*; Uk: Unknown.

Our core taxa analysis identified nine generalist virtual taxa commonly associated with citrus in both Florida and California orchards (Fig. 4) and included two *Dominikia* taxa (VTX00156, −304), three *Rhizophagus* taxa (VTX00092, −100, −248), one *Septoglomus* taxon (VTX00063), and three unknown taxa within the Glomeraceae (VTX00214, −301, −323). The remaining AMF taxa were associated with a single geographical location, and under specific management practices or disease phenotype, which could imply specialized functions and/or unique growth needs (Figs. 4–6). Therefore, we refer to these taxa as “specialist”. Several taxa from both the generalist and specialist groups were differentially abundant across geographical location (Fig. 4; Wald’s Test: *P* < 0.01), including three generalist taxa (*Rhizophagus* VTX00092, *Septoglomus* VTX00063, and unknown Glomeraceae VTX00214) and nine specialist taxa, with five from Florida (*Rhizophagus* VTX00113 and −115; *Dominikia* VTX00222 and −125; unknown Glomeraceae VTX00384) and four from California (*Dominikia* VTX00155 and −130; *Septoglomus* VTX00409 and −64). Our data also indicated that organic and conventional California orchards shared ten AMF taxa, including eight of the nine generalist taxa. One generalist taxon (unknown Glomeraceae VTX00323) was only associated with conventional orchards. In contrast, several AMF taxa were unique to organic orchards including two *Septoglomus* (VTX00064, −331), one *Rhizophagus* (VTX00105), and three *Dominikia* (VTX130, −175, and ‘Unknown’), among which *Dominikia* VTX00130 and *Septoglomus* VTX00064 were significantly enriched (Fig. 5: Wald’s Test: *P* < 0.01). In addition, AMF taxa were clearly impacted by HLB as nine VT could not be detected in trees severely affected by HLB (Fig. 6). Specifically, generalist unknown Glomeraceae VTX00323 and *Dominikia* VTX00132 were significantly depleted as HLB symptoms became more severe and could potentially serve as biological markers of tree health or HLB (Fig. 6: Wald’s Test: *P* < 0.01).

## Discussion

Here, we report the first comprehensive study of AMF diversity among US citrus orchards in different growing regions, under different management strategies, and under different levels of disease severity using modern sequencing methods and computational analysis. We found that AMF richness was similar between Florida and California and both citriculture regions shared a set of generalist taxa. However, AMF community composition varied significantly among these two distinct climatic zones with some specialist taxa being unique to a specific region. Our results indicated that organic practices supported AMF diversity with a unique set of specialist taxa not detected in conventionally farmed orchards. In addition, the worsening of disease symptoms severity in HLB impacted orchards induced a significant decrease in AMF richness accompanied with a drastic shift in community composition.

Our results indicate that citrus trees were commonly found (78%) in association with AMF as reported in other systems (1, 2) and that AMF community composition in US orchards was more diverse than initially reported. The last comprehensive AMF study in California and Florida orchards was based on spore morphology and identified the cosmopolitan genus *Glomus*, as well as the two genera *Gigaspora* and *Sclerocystis* (11). Previous studies on citrus that have used morphology and DNA sequencing-based approaches also reported that *Glomus* species were dominant AMF taxa in orchards worldwide including Brazil (12), China (13, 48) and Spain (49). In contrast, we found a broader number of AMF genera including *Dominikia, Funneliformis, Glomus, Rhizophagus, Septoglomus*, and likely additional undescribed genera within the Glomeraceae. Our results also indicated that *Dominika* and *Rhizophagus* were the most abundant genera associated with citrus roots, although the *Dominikia* group was not well supported. We achieved this fine-scale resolution of citrus-associated AMF clades by adopting a novel taxonomy assignment approach based on AMF-specific amplicons and developed by Stefani et al. (26). This disparity between our results and previous reports can be explained by the recent taxonomic revision of the Glomeromycota phylum in which the family Glomeraceae was split into several families (http://www.amf-phylogeny.com/; (37)) and many *Glomus* species were moved to different genera. Based on this fact, our results partially corroborate previous findings. Hence, the *Glomus* virtual taxa associated with citrus in China identified by Song et al. (48) based on the MaarjAM database, clustered in our analysis with *Rhizophagus*, *Glomus*, *Septoglomus* and *Dominikia*. Furthermore, *Glomus fasciculatus* and *G. constrictus*, identified by Nemec et al. (11) using spore morphology have now been renamed *Rhizophagus fasciculatus* and *Septoglomus constrictum*, respectively, both of which were clearly identified in our study. However, we did not identify taxa outside of the Glomeraceae, whereas *Gigaspora* spp. (Gigasporaceae) and *Glomus etunicatus* (now named *Claroideoglomus etunicatum*, Claroideoglomeraceae) were previously described by Nemec et al. (11). This difference may be due to our sampling size, seasonal variation (50), temporal variation (nearly 40 years between sampling events), location and edaphic properties of orchards sampled (29), and variety of citrus sampled (14), which can all affect AMF composition. Together these results suggest that the AMF communities associated with trees in citriculture across the globe are composed of a diverse set of taxa.

The AMF community in Florida and California was composed of virtual taxa that were shared in both states while others were specific to each state. The most abundant and common virtual taxa within the genera *Dominikia* (VTX00156, VTX304), *Rhizophagus* (VTX00092 and VTX00248), *Septoglomus* (VTX00063) and unidentified Glomeraceae species (VTX00214 and VTX00301), were also associated with citrus in China (48), apple in Italy (51), and barrel medic (*Medicago truncatula*) in Tunisia (52) suggesting that they are not specific to citrus or the US. *Rhizophagus fasciculatus* and *Septoglomus constrictum* have been classified as generalist AM fungi capable of colonizing a broad range of soils (8), and our data corroborate these findings. In fact, *Rhizophagus* species have been used broadly in agriculture to improve soil, promote host plant growth, and cope with diseases (53, 54). However, the nature of the interaction between AMF and its host are context dependent and may not always result in beneficial mutualistic outcomes (55, 56). Moreover, the specialist *Rhizophagus* taxa identified in Florida (VTX00113 and −115), were previously found in Tunisia (52) perhaps indicating that some environmental conditions in those geographic areas were not present in California. Additional sampling will need to be performed from citrus-producing areas to classify specialist from generalist taxa and identify those that confer beneficial functions on citrus with potential to be deployed as probiotics commercially. Overall, these data support the global distribution of ubiquitous taxa (36, 57). The community segregation in the two distinct citrus climatic zones within the continental US indicates that community composition is driven by environmental conditions and ecological requirements of AMF (58).

Our study also took advantage of modern sequencing methods to evaluate the impact of farming practices on AMF community richness and composition. Farming practices and soil characteristics have been recognized to affect soil microbial biodiversity and fertility (6, 7). AMF are major components of soil agroecosystem structure, functionality, and productivity, and low input agricultural practices (e.g., organic farming) have been shown to be conducive to AMF biodiversity, activity, and root colonization in both annual (9, 59) and perennial cropping systems (12, 51). Our results support those findings as we measured distinct AMF community profiles in organic orchards in comparison to nearby conventional orchards. In addition, several virtual taxa, including *Rhizophagus* (VTX00105) and *Septoglomus* (VTX00064), were specifically enriched in organic orchards. Exogenous application of *S. constrictum* and *R. intraradices* stimulated plant growth or productivity in several cropping systems and increased tolerance to heat and drought stress (60, 61). Soil organic matter content, pH and phosphorus availability are major factors that shape AMF community composition and assemblage (8, 29), and one can speculate that edaphic factors were affected under organic farming practices and in turn influenced the symbiotic relationship between AMF and citrus roots. Additional research will need to evaluate if *S. constrictum* and *R. intraradices* provide some fitness advantage for citrus trees under the semi-arid conditions of California’s primary citrus growing areas.

We found conclusive evidence that increased severity of HLB disease negatively impacts AMF richness, which corroborates previous findings using ITS2 amplicons (22). We used the same citrus root samples from Florida collected by Ginnan et al. (22) but sequenced the 18S rDNA in place of the ITS region, which is a more robust approach to quantify and profile the Glomeromycota (27, 28, 62). Our data confirmed a depletion of AMF richness and shifts in community composition as HLB severity worsened. AMF are obligate plant root symbionts and as such are greatly dependent on the host root carbon resources for metabolic processes. Trees expressing severe HLB symptoms, with thin canopy, small leaves, and branch dieback also show significant root collapse. These symptoms are caused by carbon sequestration in the aboveground tissues and poor belowground translocation of photoassimilates leading to root carbohydrate starvation (63) and likely results in a disturbance of the symbiosis with AMF. In parallel, the mycorrhizal decline was also coupled with a significant increase in parasitic fungi and oomycetes (i.e., *Fusarium* and *Phytophthora*) of the host rhizosphere and this enrichment of root-associated parasites was proposed to contribute to the root decline and hasten the decline of trees impacted with HLB (22). Several studies have highlighted the instrumental role of AMF in delaying disease onset or reducing symptoms against the soilborne pathogens *Fusarium* and *Phytophthora* in several pathosystems (64, 65) including citrus (24, 25). An AMF-induced disease resistance mechanism has been described as an integrated part of the symbiosis between the host and its AMF-associated root community. Hence, the host defense response can be either localized in the roots or systemic throughout the plant and can be activated either constitutively or primed upon pathogen attack (3). We propose that citriculture practices that foster AMF-citrus root symbioses may be able to sustain tree health under disease pressure and especially expand tree longevity and productivity in the context of HLB.

In conclusion, our unique and innovative approach to study rhizosphere mycobiome has greatly increased our understanding of AMF communities in citrus and perennial agroecosystems at large. A comprehensive and robust taxonomic assignment will benefit future AMF-root host interaction inquiries with major implications for improving citrus protection and yield.

## Acknowledgements

We acknowledge James Randolph’s contribution for training MengYuan Xi in molecular methods. We thank the California Citrus Research Board (Grant Numbers 5300-164 and 6100), the California Department of Food and Agriculture (Grant Number SCB16056) and National Institute of Food and Agriculture (Grant Number 2017-70016-26053) for funding support. We also acknowledge the National Science Foundation Graduate Research Fellowship Program Directorate for Biological Sciences (Grant Number NSF DGE-1326120). Any opinions, findings, and conclusions or recommendations expressed in this material are those of the author(s) and do not necessarily reflect the views of the National Science Foundation.

## Author contributions

PER, SIG, MCR, and GV designed the project. PER, NG, TD, SB collected root samples. NG, TD and SB extracted DNA from samples. MYX prepared DNA libraries. MYX, ED and VETMA performed computational and/or phylogenetic and created the figures. PER, SIG, MYX and ED wrote the manuscript. Everyone contributed significantly to the manuscript and gave final approval for publication.

